# Decline of auditory-motor speech processing in older adults with hearing loss

**DOI:** 10.1101/169235

**Authors:** Muriel TN Panouillères, Riikka Möttönen

## Abstract

Older adults often experience difficulties in understanding speech, partly because of age-related hearing loss. In young adults, activity of the left articulatory motor cortex is enhanced and it interacts with the auditory cortex via the left-hemispheric dorsal stream during speech processing. Little is known about the effect of ageing and age-related hearing loss on this auditory-motor interaction and speech processing in the articulatory motor cortex. It has been proposed that up-regulation of the motor system during speech processing could compensate for hearing loss and auditory processing deficits in older adults. Alternatively, age-related auditory deficits could reduce and distort the input from the auditory cortex to the articulatory motor cortex, suppressing recruitment of the motor system during listening to speech. The aim of the present study was to investigate the effects of ageing and age-related hearing loss on the excitability of the tongue motor cortex during listening to spoken sentences using transcranial magnetic stimulation and electromyography. Our results show that the excitability of the tongue motor cortex was facilitated during listening to speech in young and older adults with normal hearing. This facilitation was significantly reduced in older adults with hearing loss. These findings suggest a decline of auditory-motor processing of speech in adults with age-related hearing loss.

## 1. Introduction

Ageing is often associated with a progressive decline in hearing, also known as presbycusis (Frisina and Frisina, 1997; Ries, 1994; see for review: Cardin, 2016), which leads to difficulties in perceiving speech (Stewart and Wingfield, 2009; Tun et al., 2010; see for review: Humes and Dubno, 2010). However, hearing loss is not the only factor affecting speech perception deficits in ageing. Indeed, older adults with normal hearing also experience difficulties in understanding speech in adverse listening conditions compared to young individuals (Frisina and Frisina, 1997; Stewart and Wingfield, 2009; Wingfield et al., 2006). Moreover, older adults with hearing deficits do not perceive speech as well as young adults with matching hearing deficits (Dubno et al., 1984; Wingfield et al., 2006). This suggests that an interaction between hearing loss and ageing compromise speech understanding. These changes in speech perception skills are associated with alterations in the structure, function and connectivity of the auditory cortex (see for recent reviews: Cardin, 2016; Peelle and Wingfield, 2016).

In young adults, the articulatory motor cortex is involved in speech perception, together with the auditory cortex (see for reviews: Liebenthal and Möttönen, 2017; Möttönen and Watkins, 2012; Schomers and Pulvermüller, 2016; Skipper et al., 2017). Models of spoken language processing suggest that the dorsal stream serves as an auditory-motor interface. This interface maps speech sounds onto motor representations via neural projections from the superior temporal cortex to the left posterior temporo-parietal junction, the left supramarginal gyrus, the left premotor and motor cortex and the left inferior frontal gyrus (Hickok and Poeppel, 2007; Rauschecker and Scott, 2009; Scott and Johnsrude, 2003). In agreement with this model, neuroimaging studies in young adults have consistently shown activation of the left frontal speech motor network including inferior frontal gyrus (IFG), premotor cortex and primary motor cortex during speech perception (Adank, 2012; Callan et al., 2010; Du et al., 2014; Hervais-Adelman et al., 2012; Londei et al., 2010; Osnes et al., 2011; Pulvermüller et al., 2006; Skipper et al., 2005; Szenkovits et al., 2012; Wilson et al., 2004.) Moreover, accumulating evidence from Transcranial Magnetic Stimulation (TMS) studies has shown that listening to speech enhances the excitability of the articulatory regions of the primary motor cortex (Fadiga et al., 2002; Murakami et al., 2011, 2013, 2015, Nuttall et al., 2016, 2017; Watkins et al., 2003). Modulation of the activity of the articulatory motor areas using TMS influences performance in both demanding speech discrimination tasks and easy word-to-picture matching tasks (Bartoli et al., 2015; D’Ausilio et al., 2009; Meister et al., 2007; Möttönen and Watkins, 2009; Schomers et al., 2015; Smalle et al., 2015). These findings demonstrate that the articulatory motor regions play a causal role in speech processing in young adults. A recent TMS study investigated the effect of disrupting nodes of the dorsal stream on both speech perception and the articulatory motor cortex excitability (Murakami et al., 2015). TMS-induced virtual lesions in the posterior regions of the dorsal stream, including the posterior superior temporal sulcus and the sylvian parieto-temporal areas, led to speech perception deficits and reduced facilitation of the articulatory motor cortex in young adults. This shows that disruptions in auditory processing in temporal areas can result in the reduced recruitment of the articulatory motor cortex during speech perception in young adults.

Little is known about the effect of age-related hearing loss and changes in auditory processing at sub-cortical and cortical levels on speech processing in the articulatory motor system. It can be hypothesized that reduced input from the cochlear to the auditory system and deficits in auditory processing reduce the input to the articulatory motor cortex. We call this the *auditory-motor decline hypothesis*. This hypothesis is based on the evidence that, in young adults, auditory and motor areas are closely connected and interact with each other from the early stages of speech processing (see, e.g., Liebenthal and Möttönen, 2017; Skipper et al., 2017). Alternatively, it can be hypothesized that the age-related hearing loss and deficits in auditory processing lead to up-regulation of the articulatory motor system, which helps maintaining speech perception. We call this *the motor compensation hypothesis*, as it is based on the assumption that the articulatory motor system has a compensatory role. There is some evidence supporting this hypothesis. Indeed, in a recent neuroimaging study, older adults showed an enhanced activation of the frontal speech motor areas relative to young adults when listening to speech in noise (Du et al., 2016). Moreover, this increased activation correlated with speech discrimination performance as older individuals with stronger activity of the articulatory sensorimotor regions also had a better accuracy in the speech perception task. The authors proposed that the up-regulation of speech motor regions compensates for the deficient auditory processing in older adults and allows successful decoding of speech in adverse listening conditions. The older adults who participated in this study had elevated hearing thresholds relative to the young adults and six out of sixteen older adults had mild-to-moderate hearing loss at a frequency relevant for speech perception (up to 4KHz, Frisina and Frisina, 1997). It is therefore unclear whether the enhanced activation of the speech motor network was caused by ageing, hearing loss or an interaction of both factors.

The aim of the present TMS study was to determine how ageing and age-related hearing loss affect the involvement of the articulatory motor cortex during listening to spoken sentences with and without noise. TMS combined with electromyography (EMG) recordings was used to assess excitability of the tongue and hand motor cortex while young and older adults listened to sentences and non-speech control stimuli. Listening to sentences was expected to enhance excitability in the tongue, but not hand, motor cortex relative to non-speech stimuli. Older adults were considered to have hearing loss when their hearing threshold was above 25dB (Schoof and Rosen, 2014) for any of the tested frequencies between 250Hz and 4kHZ (Frisina and Frisina, 1997). First, we tested whether ageing affected the recruitment of the articulatory motor cortex during speech perception by comparing the excitability of the tongue motor cortex between young and older adults with normal hearing. Second, we assessed the differences in facilitation of the tongue motor cortex when listening to speech between older adults with normal hearing (NH) and older adults with hearing loss (HL) in order to test whether age-related hearing loss affects speech processing in the articulatory motor cortex. We had two alternative hypotheses regarding the effect of age-related hearing loss on the involvement of the articulatory motor cortex in speech perception: (1) *The motor compensation hypothesis* predicts that age-related hearing loss will increase the recruitment of the articulatory motor cortex during speech perception, which will compensate for auditory deficits. (2) *The auditory-motor decline hypothesis* predicts that age-related hearing loss will reduce the recruitment of the articulatory motor cortex during speech perception.

## 2. Material and Methods

### 2.1. Participants

A total of twenty-one young participants and twenty-four older adults were recruited for the present research. The data of three young adults were excluded because of a failure to record tongue motor evoked potential, the behavioural performance on the 0dB sentence report being below 50% accuracy or the MOCA (Montréal Cognitive Assessment) score being smaller than 26. The data of three older adults was also rejected as their score on the MOCA was below 26. Thus, we report the data of eighteen young adults (mean age: 21.9 ±2.9, range: 18-26 years; 10 females) and twenty-one older adults (mean age: 68.7 ±3.2, range: 63-74 years; 8 females). 10 older adults had a clinically normal hearing (NH) within the speech frequencies (250 Hz - 4 KHz), whereas 11 older adults had a hearing loss (HL) within the speech frequencies (250 Hz - 4 KHz). All participants were right-handed and native-English speakers with no known neurological, psychiatric or language impairment. All participants gave their written informed consent and were screened prior inclusion for contraindications to TMS. Experimental procedures conformed to the Code of Ethics of the World Medical Association (Declaration of Helsinki) and were approved by the Central University Research Ethics Committee (CUREC) at the University of Oxford (CUREC Reference: R45417/RE001).

Participants’ musical abilities were evaluated using the training sub-scale of the Goldsmiths Musical sophistication index (Müllensiefen et al., 2014), which is a self-report inventory assessing individual differences in musical sophistication. Musical abilities were assessed because it has been shown that musical training can have a beneficial effect on speech perception in noise, both in young and older adults (see for review: Alain et al., 2014). Depression was evaluated via the Beck Depression Inventory (BDI-II). Participants had minimal to mild depression (min: 0 – max: 19). We controlled for depression because it can lead to cognitive impairments (Mathews and MacLeod, 2005) which could have affected performance on some of the tasks used in the present study such as the QuickSIN and the speech report task.

### 2.2. Experimental design

All subjects received one block of single-pulse TMS to the tongue area of left primary motor cortex and one block of single-pulse TMS to the hand area of left primary motor cortex as a control site, while listening to clear speech, speech in noise, speech-correlated noise or white noise. We recorded from the tongue muscle and not from the lip muscle as we did in previous studies, because stimulating the lip area can be challenging for a number of reasons. First, lip stimulation may lead to unreliable motor-evoked potentials (MEPs - (Möttönen and Watkins, 2009; Panouillères et al., 2018; Rogers et al., 2014; Swaminathan et al., 2013). Second, the intensity of the stimulation required to induce an MEP in the relaxed lip muscle is usually quite high (ranging from 60 to 70% of the maximum intensity output of the Magstim 200). This level of intensity can lead to discomfort that is not always tolerated by participants (Adank et al., 2017; Möttönen and Watkins, 2009; Rogers et al., 2014). Thus, to minimise the exclusion rate based on unreliable MEPs and discomfort, we targeted the motor representation of the tongue. The order of the hand and tongue blocks was counterbalanced across participants. Following the two TMS blocks, participants completed a short task in which they verbally reported sentences in clear speech and sentences in noise. Their hearing thresholds and speech-in-noise abilities were then assessed using respectively pure-tone audiometry and QuickSIN (Etymothics - (Killion et al., 2004)). Finally, participants’ depression, cognitive abilities and musical training were assessed using various questionnaires (see above) and the participants performed a working-memory task.

### 2.3. Electromyography

Electromyography (EMG) activity from the right first dorsal interosseous (FDI) muscle was recorded by placing a pair of electrodes (22 × 30 mm ARBO neonatal electrocardiogram electrodes) on the belly and tendon of the muscle. In order to record MEPs from tongue muscles, two disposable EEG cup electrodes (Unimed, UK) were mounted to a nose-clip (Cressi, Italy). Participants placed this device into their mouth so that the two electrodes were above and below the right side of the tongue. The ground electrode was attached to the right wrist. The raw EMG signal was amplified (gain: 1000), band-pass filtered (1-1000Hz) and sampled (5000Hz) via a CED 1902 four-channel amplifier, a CED 1401 analog-to-digital converter and a computer running Spike2 (Cambridge Electronic Design). This EMG signals were stored on the computer for off-line analysis.

### 2.4. Transcranial Magnetic Stimulation

All TMS pulses were monophasic, generated by Magstim 200 (Magstim, Whitland, UK) and delivered through a 70-mm figure of eight coil. The position of the coil over the left motor cortex was adjusted until a robust motor-evoked potential (MEP) was observed in the contralateral target muscle (either hand or tongue). The intensity of the stimulation was set as the lowest intensity consistently eliciting reliable MEPs in the resting muscle. The mean intensity used for the tongue muscle was of 59.7% (±SE:2.4%) for the young participants and of 61.8% (±SE: 2.3%) for the older adults. For the hand stimulation, the averaged intensity was of 51.5% (±SE: 2.9%) and of 55.2 (±SE: 2.7%) for the young and the older adults, respectively. The stimulation intensity significantly differed between the stimulation sites (F[1,37]=13.30, p<0.001) but not between the age groups (group effect: F[1,37]<1, p=0.34; group × site interaction: F[1,37]<1, p=0.68).

### 2.5. Speech stimuli

The stimuli used in the present study were selected from a set of sentences used in previous studies (Davis et al., 2011; Rodd et al., 2005). The set comprised 100 declarative sentences between 6 and 13 words in length that were semantically coherent. All 100 sentences (1.2 to 3.5 seconds in duration, speech rate 238 words/minute) were produced by a male speaker of British English and digitized at a sampling rate of 44.1 KHz. For the speech in noise condition, the sentences were degraded by adding speech-spectrum signal-correlated noise (SCN) at a signal-to-noise ratio (SNR) of 0dB using Praat software (Davis et al., 2011). Fourteen pure SCN stimuli and fourteen white noise (WN) stimuli were also used in the present study as control stimuli.

### 2.6. Design of the TMS blocks

Participants sat in front of a computer presenting the stimuli using Presentation^®^ software (Neurobehavioral Systems, Inc., Berkeley, CA, USA). Audio stimuli were presented to the participants through insert earphones (Etymotic, Elk Grove Village, IL, USA). For each TMS site (hand and tongue), participants were exposed to 20 clear sentences, 20 sentences in noise, 30 SCN stimuli and 30 WN stimuli. For each block, the order of these 100 stimuli was randomized. Altogether, the two TMS blocks (tongue and hand) included 40 clear sentences, 40 sentences in noise, 60 SCN stimuli and 60 WN stimuli. Each sentence was presented only once. Participants were instructed to listen to the stimuli while keeping both their tongue and hand muscles relaxed. For each stimulus (clear speech, speech in noise, SCN or WN), a single-pulse of TMS was delivered to elicit an MEP. For the sentence stimuli, it was delivered 150 ms after the onset of the final content word, as in our previous study (Panouillères et al., 2018). This was chosen as a reliable way of matching the point at which TMS was delivered across sentences, as the final content word was likely to be the most predictable. For the WN and SCN stimuli, the pulse was delivered close to the end of the stimuli, matching the timing of the pulses for the sentence stimuli. The average inter-trial interval was set at 5s (range: 4.02s - 5.99s). Each block lasted ~7 minutes and within each block, a short break occurred every 33-34 trials.

### 2.7. Speech report task

After the completion of the two TMS blocks, participants completed a speech report task to assess their ability to perceive speech. Participants were asked to listen to 10 sentences in clear speech and to 10 sentences in noise (SNR of 0dB), randomly presented, and after each of them, they were asked to repeat the sentence. The accuracy of the response was evaluated during the task by the experimenter, and off-line using the audio-recordings of the responses. Once the participants were finished repeating a sentence, they could move on to the next trial by pressing the space bar.

### 2.8. Pure-tone audiometry and QuickSIN

Pure-tone audiometric hearing thresholds were assessed using a diagnostic audiometer (Avant A2D+, MedRX International, Germany) across 250 Hz, 500 Hz, 1000 Hz, 2000 Hz, 4000 Hz and 8000 Hz for both ears. Older participants were divided into two sub-groups based on their hearing thresholds upon the range for speech perception from 250 Hz to 4 KHz (Frisina and Frisina, 1997). The normal hearing (NH) older adult group had normal thresholds ≤ 25dB from 250 Hz to 4 KHz, while the older adult group with hearing loss (HL) had mild-to-moderate hearing impairment, with thresholds > 25dB on these same frequencies. Pure-tone average (PTA) audiometric thresholds were computed across frequencies from 500 Hz to 4 KHz for each ear.

The Quick Speech-In-Noise (QuickSIN) test was used to assess speech perception in noise (Killion et al., 2004). This test is composed of lists of low-context sentences spoken by a female talker presented in a four-talker babble background. The test was presented binaurally through the Avant A2D+ audiometer, with the speech levels set to 70 dB. Each list contained 6 sentences with a SNR decreasing from 25 dB to 0dB in 5dB steps. Following each sentence, participants were instructed to repeat to the experimenter as many words as possible. Each sentence was scored based on the correct recognition of five “target” words (e.g. <u>Crouch before</u> you <u>jump</u> or <u>miss</u> the <u>mark)</u>. The test score, called SNR loss, represents the SNR required for the listener to repeat 50% of the words correctly. It was calculated for each list as the difference between 25.5 and the total number of words correctly reported for that list. One practice list and four test lists were presented per participant; the order of the test lists being counterbalanced across participants. The average SNR loss across the four test lists was calculated to obtain the final score.

### 2.9. Working-Memory reading span task

We assessed participants’ working memory because it has been shown to be linked with speech perception in noise (Akeroyd, 2008). We administered the shortened version of the reading span task, which has been verified to retain the psychometric properties of the longer version (Foster et al., 2015; Oswald et al., 2015). The task was automated, presenting stimuli on screen and collecting the responses via mouse click (task downloaded from: http://englelab.gatech.edu/). Participants were presented with a set of sentences of ~10 to 15 words and they had to decide whether the semantic content of the sentence was correct or not. After each sentence, participants were presented with a letter for recall at the end of the set. Set sizes ranged from 4 to 6, with two administrations for each set size. Participants practiced each task (remembering the letters and judging the semantic content of the sentence) separately and then two trials of the two tasks combined before starting with the main test. Participants’ score was obtained by averaging the proportions of correctly recalled elements in each trial.

### 2.10. MEPs analysis

MEPs were analysed on a trial-by-trial basis using in-house software written in Matlab (Mathworks Inc, Natick, USA). Maximal and minimal peaks of the MEPs were automatically detected using a fixed time window following the TMS pulse: [15-40ms] for the hand and [12-35ms] for the tongue. The detection was checked manually by the experimenter. The absolute value of the background muscle activity was averaged within a 100-ms window preceding the TMS pulse and trials with a mean absolute value of background muscle activity higher than 2 standard-deviations of the average for each TMS block were excluded. Outliers MEP with values above or below 2 standard-deviations of the mean for each experimental condition (clear speech, speech in noise, SCN and WN) were removed. We calculated MEP z-scores for each experimental condition relative to the WN in each participant, separately for the hand and the tongue blocks.

### 2.11. Statistical analysis

Statistical analyses were performed with the SPSS Statistics software package (IBM, Armonk NY, USA). Participants’ level of depression, years of education, MOCA, musical training, working memory, PTA and SNR loss (QuickSIN) were compared with separate one-way ANOVAs with the between-subjects factor group (young vs NH vs HL). Performance on the speech report task was submitted to a mixed design ANOVA with the between-subject factor group (young vs NH vs HL) and the within-subject factor noise level (clear speech vs speech in noise). To investigate the effect of age on the recruitment of the primary motor cortex on speech perception, separate ANOVAs for the hand and tongue stimulation were performed on the MEP z-scores with the between-subject factor group (young vs NH) and the within-subject factor stimulus (speech in noise vs clear speech vs SCN). Similar ANOVAs were carried out to compare the MEP z-scores between NH and HL groups. Significant main effects or interactions in the ANOVAs were followed by bootstrapping for independent samples t-test (1000 sampling), and corrected for multiple comparisons (Bonferroni).

Correlational analyses were run separately for the young and older adults and for the hand and tongue stimulation between the MEP z-scores (averaged for the speech in noise and clear speech stimuli) and Participants’ PTA. Similar correlations were performed between the z-scores for the SCN stimuli and the PTA. Bayesian analyses were performed with JASP software (http://jasp-stats.org/). The main purpose of the Bayesian analyses was to assess the amount of support for the non-significant results, which are often misinterpreted (see e.g., Aczel et al., 2018). Significance was set at p<0.05.

## 3. Results

### 3.1. Participants’ characteristics

The characteristics of the three groups of participants (young adults, older adults with NH and older adults with HL) are presented in Table 1. All three groups were matched in total number of years of education and the level of depression (one-way ANOVA: F[2,36]<1, p>0.39). Only older adults with normal cognitive abilities were included in the present study (MOCA≥26). However, older adults with HL had slightly lower cognitive abilities than the young adults and the older adults with NH (F[2,36]=5.18, p<0.05; Bonferroni post-hoc tests: young vs HL: p<0.01 and NH vs HL: p<0.05 respectively). The young adults had received overall more musical training than the NH and HL older adults group (F[2,36]=4.90, p<0.05; Bonferroni post-hoc tests: young vs NH: p<0.05 and young vs HL: p<0.01 respectively). Although there was a tendency for the older adults to have a lower score in the reading span task assessing working memory, this difference was not statistically significant (F[2,36]=2.22, p=0.12).

**Table 1:** Participants characteristics. MOCA: Montreal Cognitive Assessment; BDI-II: Beck Depression Inventory II; WM: Working-Memory. Errors are standard-error of the mean.

|  | Age (years) | Years of education | Musical training | MOCA | BDI-II | WM score (%) |
| --- | --- | --- | --- | --- | --- | --- |
| Young | 21.9 ±0.7 | 16.8 ±0.4 | 20.7 ±3.5 | 28.8 ±0.3 | 2.6 ±0.9 | 66.7 ± 5.7 |
| Older NH | 68.0 ±1.2 | 16.1 ±0.9 | 7.5 ±1.7 | 28.4± 0.3 | 3.8 ±1.8 | 50.0 ±6.6 |
| Older HL | 69.4 ±0.7 | 15.2 ±1.1 | 9.8 ±3.5 | 27.6 ±0.3 | 2.9 ±1.5 | 47.0 ±10.7 |

### 3.2. Participants’ hearing abilities and speech perception performance

The results of the pure tone audiometry averaged across both ears for all the tested frequencies are plotted in Fig. 1A. The young adults had normal hearing thresholds at all tested frequencies. For the older adults with NH, pure-tone thresholds were in the normal range in the speech frequencies range (250 Hz to 4 KHz), but these participants had moderate hearing impairments at the highest frequency (8 KHz). The hearing thresholds of the older adults with HL were in the normal range from 250 Hz to 2 KHz (below 25 dB), but they were in the moderate and moderately severe hearing loss range for the 4KHz and 8KHz frequencies, respectively. The pure tone averages (PTA) from 500 Hz to 4 KHz (Fig. 1B) differed significantly between the groups (F[2,36]=45.60, p<0.001). Indeed, both groups of older adults, including the group with normal hearing, had higher PTA than the young adults (Bonferroni post-hoc tests, p<0.01). As expected, the older adults with HL had higher PTA than the older adults with NH (Bonferroni post-hoc test, p<0.01).

**Figure 1:**
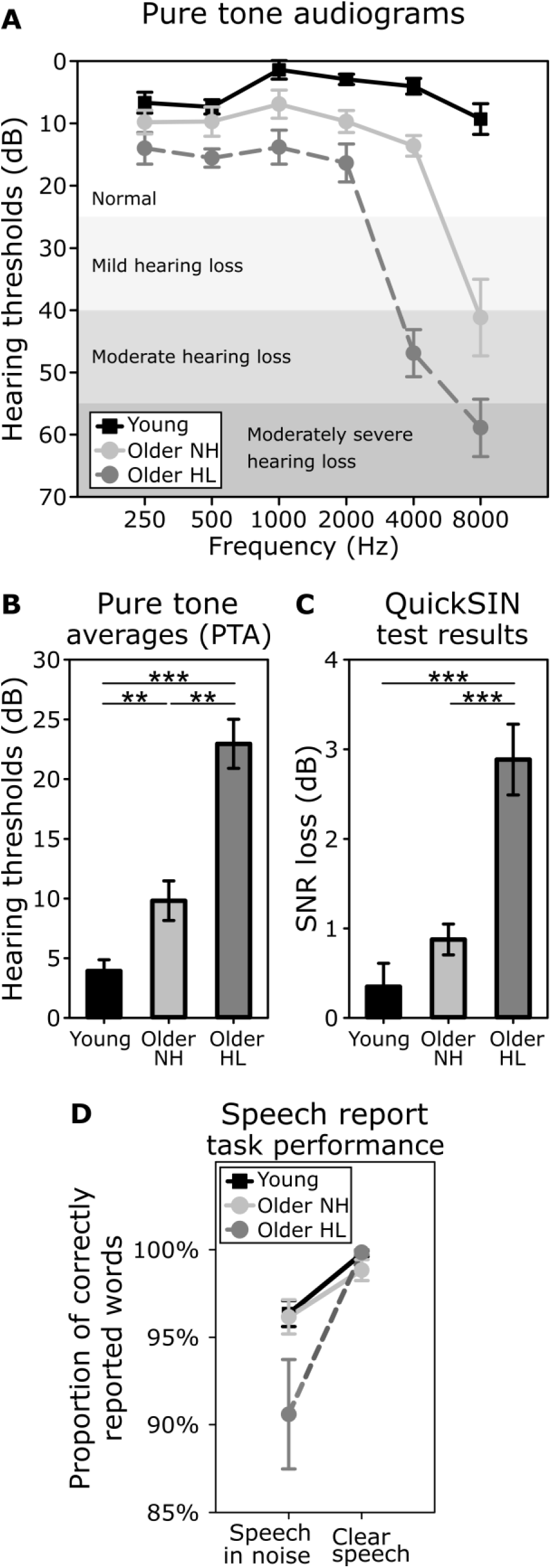
Hearing and speech perception abilities. (A) Pure tone audiometry results. The hearing thresholds are represented averaged for both ears. (B) Pure tone averages (in dB) across the frequencies 500Hz to 4 KHz. (C) QuickSIN results presented as Signal-to-Noise Ratio (SNR) loss. (D) Proportion of correctly reported words in the speech report task performances. Data are presented in black for the young adults, in light grey for the older adults with NH and in dark grey for the older adults with HL. Error-bars are standard-error of the mean. Significant differences between groups are indicated by: ** p<0.01 and *** p<0.001 (Bonferroni post-hoc tests).

With the QuickSIN test, we measured the SNR loss of each participant, which is defined as the dB increase in SNR required to understand speech in multi-babble noise relative to someone with normal hearing (Fig. 1C). Older adults with HL had greater SNR loss scores than young adults and older adults with NH (Group effect: F[2,36]=20.02, p<0.001; Bonferroni post-hoc tests: p<0.001). The normal range for this test is considered to be below 2dB of SNR loss (Killion et al., 2004). The SNR loss scores of more than half of the older adults with HL (7/11) were equal or greater than 2dB, whereas the SNR loss scores of none of the older adults with NH (0/10) were equal or greater than 2dB. The difference in SNR loss scores between older adults with NH and young adults was non-significant (Bonferroni post-hoc test: p=0.10). This non-significant effect should be however interpreted with caution, because Bayesian analyses showed that the support for the null hypothesis (i.e., no effect of age) was anecdotal (BF_01_=1.34).

Speech perception skills were also tested using a speech report task, in which participants listened to clear sentences and sentences in speech-correlated noise at SNR of 0dB. After each sentence, participants had to verbally report as many words as possible. The sentences were similar to the ones the participants passively listened to during the TMS part of the experiment. The proportion of correctly reported words was reduced for the 0dB stimuli compared to the clear stimuli (Fig. 1D – Stimulus effect: F[1,36]=24.03, p<0.001). The performance of the groups of participants differed for sentences in noise, but not for clear sentences (Group effect: F[2,36]= 3.12, p=0.06; Stimulus × Group interaction: F[2,36]=3.75, p<0.05) as the older adults with HL reported the sentences in noise slightly less accurately than the other groups (Fig. 1D).

### 3.3. No effect of age on the excitability of motor cortex when listening to speech

To investigate the effect of ageing on the excitability of the primary motor cortex when listening to speech and non-speech sounds, we compared the MEP z-scores between the young adults and the older adults with NH, separately for the hand and tongue stimulation (Fig. 2 A and 2B). The facilitation of the hand motor cortex was similar for both sentences and SCN for both age groups (Stimulus effect: F[2,42]<1, p=0.52; Group effect: F[1,26]=1.52, p=0.23; Stimulus × Group interaction: F[2,42]<1, p=0.96), suggesting that the modulation of hand motor excitability was not specific to speech (Fig. 2B). In contrast, the facilitation of the tongue motor excitability was stronger for the sentences than for the SCN (Fig 2A – Stimulus effect: F[2,52]=7.29, p<0.01; Bonferroni post-hoc tests: speech in noise versus SCN: p<0.05; clear speech vs SCN: p<0.01, speech in noise versus clear speech: p = 1). Finally, there was no significant difference in the tongue motor excitability between young adults and older adults with NH (Group effect: F[1,26]=1.04, p=0.32; Stimulus × Group interaction: F[2,52]=2.21, p=0.12). This should be however interpreted with caution because Bayesian analyses showed that the support for the null hypothesis is anecdotal (BF01 = 1.51 for Stimulus × Group interaction).

**Figure 2:**
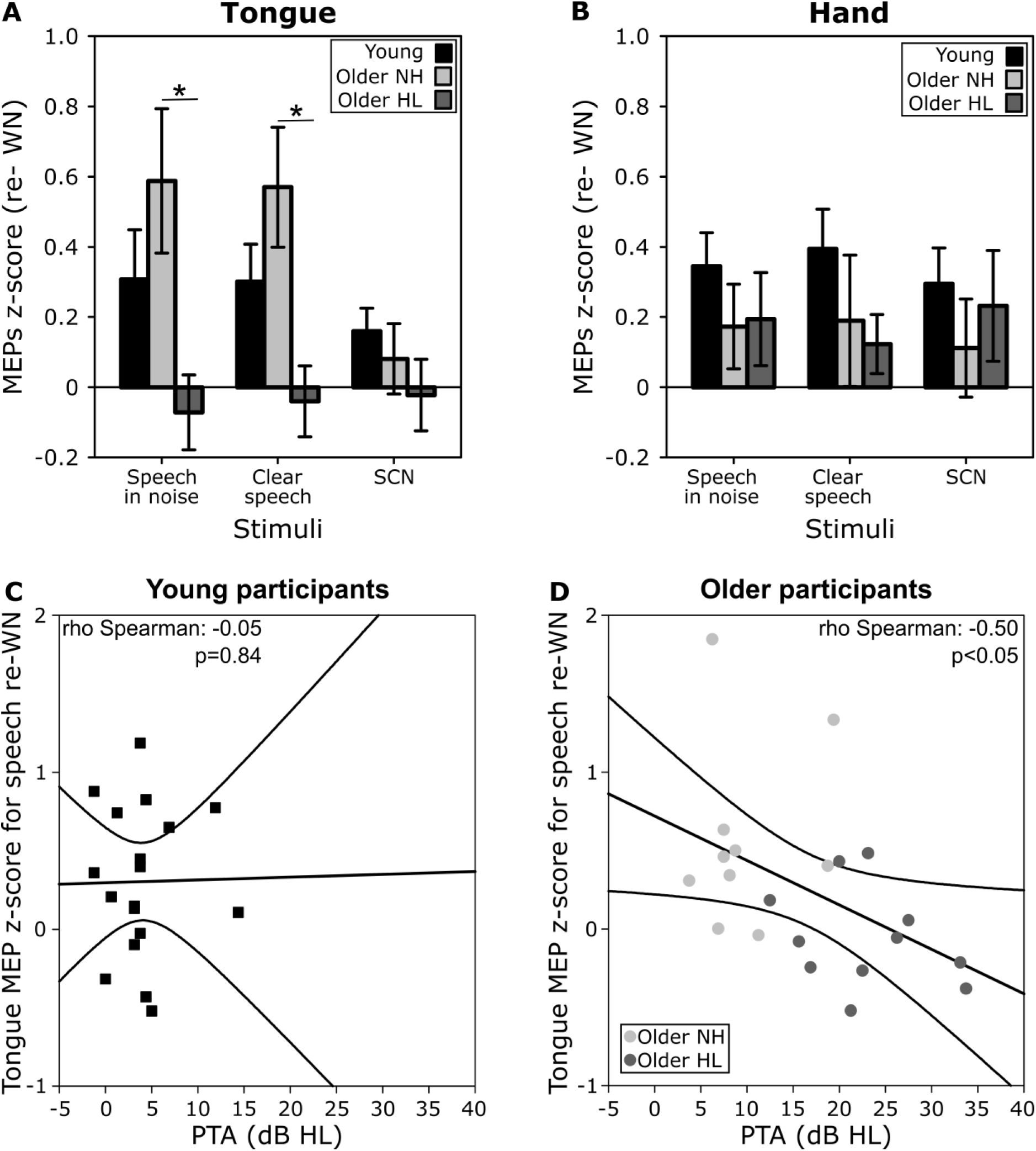
MEP z-scores during speech perception. The MEPs elicited during the perception of clear speech, speech in noise and Speech-correlated noise (SCN) are represented as z-scores normalised to WN for the tongue (A) and hand (B) conditions. Error bars are standard error of the mean. Significant differences between groups are indicated by: * p<0.05 (Bonferroni post-hoc tests). Correlations between the averaged tongue MEP z-scores for the clear speech and speech in noise condition and pure tone averages (PTA) are represented for the young participants (C) and the older participants (D). Data are presented in black for the young adults, in light grey for the older adults with NH and in dark grey for the older adults with HL. Correlations are represented with the confidence interval.

### 3.4. Age-related hearing loss reduces the excitability of the tongue motor cortex during listening to speech

To investigate the effect of age-related hearing loss on the motor excitability, when listening to speech and non-speech sounds, we compared the MEP z-scores between the older adults with HL and the older adults with NH, separately for the hand and tongue stimulation (Fig. 2A and 2B). The facilitation of the hand motor excitability was similar for both sentences and SCN and for the two groups of older adults (Fig. 2B - Stimulus effect: F[2,28]<1, p=0.93; Group effect: F[1,19]<1, p=0.88; Stimulus × Group interaction: F[2,28]<1, p=0.59). The facilitation of tongue motor cortex was reduced in older adults with HL compared to the older adults with NH as a function of the stimuli (Fig 2A – Stimulus effect: F[2,27]=4.92, p<0.05; Group effect: F[1,19]=7.90, p<0.05; Stimulus × Group interaction: F[2,27]=6.47, p<0.01). The Bayesian analyses showed that the support for this effect was strong (BF_10_ = 10.58 for Stimulus × Group interaction). More specifically, the excitability of the tongue motor cortex was reduced in the HL group relative to the NH group for the clear sentences and the sentences in noise (Bonferroni post-hoc tests: speech in noise and clear speech: p<0.05, SCN: p=0.47). Because the z-scores were normalised to the WN MEPs, we assessed whether there could be any difference between older adults with HL and older adults with NH in this baseline condition. The WN MEPs were similar between the two groups (independent samples t-test: t[19]=- 1.09, p=0.29), showing that the decreased excitability of the tongue was not a general decreased of excitability but was specific to listening to speech. We finally assessed whether the MEPs may differ from the WN baseline in the older with HL by comparing the z-scores for the tongue session with 0. We did not find any differences from 0 (t[10]=[-0.67; −0.22], p>0.52), confirming the lack of facilitation of the tongue motor cortex when listening to speech and SCN relative to WN. Thus, our results show that listening to speech does not enhance the excitability of the tongue motor cortex in adults with age-related hearing loss.

### 3.5. Correlation between tongue excitability when listening to speech and hearing thresholds

To further explore the relationship between motor excitability when listening to speech and hearing thresholds, we ran correlational analyses for the young and older adults separately (Figure 2C and 2D). The tongue MEP z-scores did not show any significant correlation with hearing thresholds in the young adult group (Fig. 2C - Spearman rho=-0.05, p=0.84). However, there was a significant negative correlation between tongue excitability when listening to speech and hearing thresholds in the older adults group (Fig. 2D - Spearman rho=-0.50, p<0.05). There was no significant correlation between hearing thresholds and tongue MEP z-scores when listening to SCN. There was no significant correlation between hand MEP z-scores and hearing thresholds in either age group (Spearman |rho|<0.16, p>0.47). Thus, the older adults with the least sensitive hearing-characterised by high PTA – had the lowest tongue motor excitability when listening to speech (relative to WN baseline).

## 4. Discussion

In this study, we demonstrate that older adults with HL at speech frequencies (up to 4kHz) show reduced recruitment of the articulatory motor cortex during listening to speech relative to older adults with NH. The older adults with HL were also impaired in speech perception in noise. The present findings suggest that reduced auditory input from the cochlear to the auditory system result in reduced recruitment of the articulatory motor system in speech processing in older adults and support the auditory-motor decline hypothesis.

The sample of older adults tested in the present study was divided into two subgroups based on whether their hearing thresholds within the speech frequency range (250 Hz to 4 KHz) was in the normal range (≤ 25dB). The group of older adults with HL had strong difficulties in perceiving speech when simultaneously presented with a multi-babble noise (multiple SNR levels, QuickSIN task) and mild difficulties for speech presented in speech-correlated noise (fixed SNR: 0 dB, speech report task). This reduced ability to understand speech in older adults with HL is in agreement with previous reports (Dubno et al., 1984; Stewart and Wingfield, 2009; Tun et al., 2010; Wingfield et al., 2006). The performance of all older adults with NH was within the normal range in the QuickSIN test and they performed similarly to young adults in both speech-in-noise tasks (QuickSIN and word report), suggesting that the skill to understand speech in noise was preserved in the group of older adults with NH in our study. However, the sample of older adults with NH was small (N=10) in our study and the Bayesian analysis showed that the evidence for the null hypothesis (i.e. no effect of age) was anecdotal in the QuickSIN test. Thus, the non-significant difference between older adults with NH and young adults should be interpreted with caution. Some previous studies have demonstrated that ageing (without hearing loss) can lead to a reduced ability to perceive speech in challenging listening conditions (Dubno et al., 1984; Füllgrabe et al., 2015; Goossens et al., 2017; Jin et al., 2014; Rajan and Cainer, 2008; Schoof and Rosen, 2014; Wong et al., 2009), whereas other studies have found no differences between young and older adults with normal hearing (Eckert et al., 2008; Frisina and Frisina, 1997; Schoof and Rosen, 2014).

In addition to hearing ability, cognitive functions and working memory are crucial for comprehending speech (Schneider et al., 2002). Moreover, musical training can have a beneficial effect on perceptual and cognitive skills, including speech perception in noise, both in young and older adults (see for review: Alain et al., 2014). Therefore, we assessed the musical training, cognitive skills and working memory performance in all participants. Young adults had more musical training and better working memory performance than both groups of older adults. However, older adults with NH performed equally with the young adults in the speech-in-noise tasks. This suggests that none of these factors contributed to speech understanding in the present study, probably because of the low level of difficulty of the tasks. Despite selecting participants based on their normal cognitive abilities (MOCA ≥26), older adults with HL had decreased cognitive functions compared to the other two groups of participants. It has been reported that hearing impairment is independently associated with poorer cognitive performance (Lin, 2011; Lin et al., 2011; Tay et al., 2006) and accelerates the rate of cognitive decline (Lin et al., 2013). The hearing loss of older adults in the present study was mild compared to the aforementioned studies (estimated based on mean PTA for 0.5 to 4 KHz above normal) but was still associated with reduced cognitive performance. This suggests that even very mild hearing modifications could be associated with subtle modification of cognitive functions. Although it is possible that the speech perception deficits of the adults with HL could partly be due to these slightly reduced cognitive abilities in the current study, we believe that this effect is minimal compared to the effect of hearing loss.

In order to measure excitability of articulatory motor cortex, we measured MEPs from the tongue while participants passively listened to spoken sentences and non-speech signals. As previously demonstrated in young adults (Murakami et al., 2011, 2015; Panouillères et al., 2018; Watkins et al., 2003), we found that listening to speech enhanced the excitability of the articulatory motor cortex relative to non-speech signals (speech-correlated noise and white noise) in young and older adults with normal hearing. Despite recent studies suggesting that the articulatory motor cortex would be more facilitated in challenging listening conditions (Murakami et al., 2011; Nuttall et al., 2016, 2017), we found that the excitability of the articulatory motor cortex was facilitated equally by clear sentences and sentences in noise, replicating the results from our previous study (Panouillères et al., 2018). Finally, we also determined that the facilitation of the motor cortex for speech stimuli relative to non-speech stimuli is specific to the articulators and no other effectors, such as the hand (Panouillères et al., 2018; Watkins et al., 2003). Thus, our results confirm previous findings about the recruitment of the articulatory motor cortex in speech perception in young adults and extend them to older adults with normal hearing.

The facilitation of the tongue motor cortex when listening to speech did not differ between young and older adults with normal hearing, suggesting that ageing itself does not modify the recruitment of the articulatory motor cortex during speech processing. It should be however noted that there was a non-significant trend towards older adults showing stronger facilitation than young adults in line with the compensation hypothesis (Du et al., 2016). The non-significant effect of age should be interpreted with caution, because the number of older participants with normal hearing was small (N=10) and the Bayesian analysis showed only anecdotal support for the null effect. Therefore, future studies with larger sample size are needed to investigate whether aging (without hearing loss) can enhance the recruitment of the articulatory motor cortex.

We found a significant correlation between hearing sensitivities and facilitation of the tongue motor cortex during speech perception in older adults, but not in young adults. Two previous TMS studies in young adults with normal hearing found a relationship between hearing abilities and facilitation of the articulatory motor cortex during speech perception (Nuttall et al., 2016, 2017). In these studies, young adults received single-pulse TMS to elicit lip MEPs while they listened to clear speech or speech in challenging conditions. The challenging conditions consisted of spoken syllables that were either presented in background noise or pronounced while a depressor was applied onto the tongue of the speaker. The results showed that there is a stronger facilitation of the lip motor cortex when individuals listened to speech in challenging conditions than to clear speech. Correlational analyses were then performed to evaluate the relationship between hearing abilities (measured as PTA) and the excitability of the lip motor cortex. In one study (Nuttall et al., 2016), there was a greater motor facilitation when listening to the challenging condition than to clear speech for individuals with less sensitive hearing, while there was an opposite finding in the other study (Nuttall et al., 2017). The authors suggested that the type of perturbation affected the direction of the correlation. Because Nuttall et al. did not use a non-speech baseline; the direct comparison with the current findings is difficult. Nevertheless, in the present study, no significant correlation was found between hearing abilities and the facilitation of the tongue motor cortex during listening to clear and noisy sentences (relative to non-speech baseline) in the young adult group. Thus, our results do not support the earlier studies suggesting that small individual differences in hearing abilities would modulate the recruitment of the articulatory motor cortex in speech perception in young adults.

In the current study, we measured motor excitability while participants listened to sentences, because we were interested in speech perception in ecologically valid conditions. The SNR level was manipulated by adding speech-correlated noise to the speech signal, producing energetic masking of the speech. Future studies are needed to determine whether some the discrepancies with earlier studies (e.g., Du et al., 2016; Nuttall et al., 2017) may be explained by differences in speech material (syllables vs. sentences) and/or the type of noise (informational vs. energetic masking).

Our main finding was that, in older adults, the hearing threshold correlated negatively with the level of motor excitability in the tongue motor cortex during listening to speech. In older adults with HL, the facilitation of the tongue motor cortex when listening to speech was reduced compared to the older adults with NH. In fact, no speech-induced motor facilitation was observed in older adults with HL. This suggests that age-related hearing loss is associated with a reduced recruitment of the articulatory motor cortex during speech processing. This finding disagrees with the motor compensation hypothesis and the results from the previous functional Magnetic Resonance Imaging (fMRI) study by Du et al (2016). In this previous study, the authors found, in a mixed group of older adults (with and without hearing loss), enhanced activation of the frontal speech motor network (relative to young adults), which correlated positively with speech discrimination performance. However, because the syllable discrimination task used in this study included a button press with the right hand on each trial, the left frontal activations could also reflect the button press, the decision related processes or the responses selection/preparation (see: Schomers and Pulvermüller, 2016 for discussion of this point). Interestingly, some neuroimaging studies have demonstrated increased activity in the prefrontal regions associated with cognitive control when older adults process speech under challenging listening conditions (Erb and Obleser, 2013; Peelle et al., 2010; Tyler et al., 2010; Wong et al., 2009). Furthermore, the activity in frontal areas and the grey matter volume of frontal areas was associated with better speech perception performance (Tyler et al., 2010; Vaden et al., 2015; Wong et al., 2009, 2010). This increased activation of frontal areas could reflect a compensatory strategy of the ageing brain to counteract the decline in sensory input. Thus, the frontal compensatory mechanisms during speech perception may be cognitive rather than motoric.

Our results are in agreement with the auditory-motor decline hypothesis. They suggest that the structural and functional changes along the auditory pathway from the cochlear to the auditory cortex that are associated with age-related hearing loss reduce the interactions with the articulatory motor cortex via the dorsal stream, leading to a reduced recruitment of the articulatory motor cortex during speech perception. Indeed, aging reduces the temporal precision of speech processing in the brain stem (Anderson et al., 2012; Bidelman et al., 2014) and modulates entrainment of low-frequency oscillations to speech signals in the auditory cortex (Presacco et al., 2016). Moreover, age-related hearing loss is associated with reduced grey matter volume and decreased activation of the auditory cortex (Eckert et al., 2012; Harris et al., 2009; Husain et al., 2011; Lin et al., 2014; Peelle et al., 2011; Yang et al., 2014). Future studies are needed to investigate how the changes along the auditory pathway lead to reduced auditory-motor interactions during listening to speech and whether this reduction in auditory-motor speech processing contributes to speech perception difficulties in older adults with hearing loss.

To conclude, this study provides novel evidence that the age-related hearing loss reduces the recruitment of the articulatory motor cortex during speech perception. Young and older adults with NH showed facilitation of the tongue motor cortex during listening to speech, whereas older adults with HL showed difficulties in speech comprehension in noise and reduced facilitation of the tongue motor cortex during speech perception. These findings demonstrate that age-related hearing loss is associated with impaired auditory-motor processing in the left dorsal stream, resulting in the disengagement of the articulatory motor cortex during speech perception.

## Acknowledgements

We thank Dr. Matt Davis and Dr. Ingrid Johnsrude for the stimulus material. We also thank Dr Jennifer Chesters, Dr Saloni Krishnan, Dr Daniel Lametti, Charlie Wiltshire and Atakan Acar for assisting with the TMS and Daniel Drew for his invaluable help in recruiting the older adults. We are grateful to the participants for contributing with their time and effort to this study.

The authors declare that there is no conflict of interests. This work was supported by the Medical Research Council, UK (G1000566).

